# Energy dissipation via the glycerol phosphate shuttle: coupling glycolysis to mitochondrial thermogenesis in *Drosophila melanogaster* flight muscle

**DOI:** 10.1101/2025.10.24.684441

**Authors:** Johannes Lerchner, Yan A. Reis, Thalita Lucio, Marcus F. Oliveira

**Affiliations:** Institute of Physical Chemistry, TU Bergakademie Freiberg, 09599 Freiberg, Germany; Laboratório de Bioquímica de Resposta ao Estresse, Instituto de Bioquímica Médica Leopoldo de Meis, Universidade Federal do Rio de Janeiro, Cidade Universitária, Rio de Janeiro, RJ, Brazil

**Keywords:** mitochondria, bioenergetics, oxidative phosphorylation, glycolysis, energy metabolism, thermogenesis, heat, efficiency, glycerol phosphate shuttle

## Abstract

Mitochondria are the central hubs of energy metabolism, integrating carbohydrate, lipid, and amino acid oxidation to produce ATP through the tricarboxylic acid cycle and oxidative phosphorylation. These organelles also regulate energy homeostasis via redox signaling and substrate exchange between cellular compartments. Mitochondrial redox shuttles maintain the balance between cytosolic and mitochondrial NAD(P)H pools by transferring reducing equivalents across membranes. Among these, the glycerol-phosphate shuttle (GPSh) connects glycolysis with mitochondrial oxygen consumption, regenerating cytosolic NAD+ while transferring electrons into the electron transport system. Although GPSh ensures continuous glycolytic flux, its lower energy yield may favor heat dissipation over ATP synthesis. In the present work, we investigated how substrate utilization shapes energy coupling and thermogenesis in *Drosophila melanogaster* flight muscle using chip calorimetry. Calorimetric assays revealed that G3P oxidation generated significantly more heat than complex I substrates, demonstrating its high thermogenic potential. When glucose was supplied to intact flight muscles, inhibition of the GPSh with iGP1 decreased heat generation by ∼85%, highlighting its importance for cytosolic NAD□turnover and glycolytic flux. In summary, GPSh serves as a key mechanism for sustaining NAD+ regeneration in glycolysis but operates with low energy efficiency, leading to increased heat production in highly metabolically active tissues such as insect flight muscle.

## Introduction

Flight represents one of the most remarkable evolutionary innovations in insects, underpinning their ability to disperse broadly, evade predators, and secure mates effectively. This remarkable capability hinges on the extraordinary energy demands of sustained flight. Insect flight muscles rank among the most metabolically active tissues known, exhibiting extraordinary adaptations for ATP production since the energy requirements to sustain flight are immense (Sacktor and Cochran 1958; Mesquita et al. 2021). Central to this capacity are the mitochondria within flight muscle cells, which due to their exceptional structural and functional specialization, support extremely high rates of oxidative phosphorylation (OXPHOS). A hallmark of these mitochondria is their densely packed inner membrane, providing an expanded surface area that correlates with respiratory capacity and ATP output (Sohal et al. 1972; Suarez et al. 2000; Gonçalves et al. 2009). Beyond energy provision, flight muscles also generate substantial heat, a feature critical to pre-flight thermogenesis that enables cold-adapted insects to safely initiate flight (Masson et al. 2017).

The glycerol-3-phosphate shuttle (GPSh) is an integral mechanism of the electron transport system (ETS) with direct contributions to ATP synthesis and heat production (Mráček et al. 2013). GPSh operates via two complementary enzymes: the cytosolic NAD□-dependent glycerol-3-phosphate dehydrogenase (cG3PDH, coded by the *Gpd1* gene), that reduces dihydroxyacetone phosphate (DHAP) to glycerol-3-phosphate (G3P) coupled to NADH oxidation to NAD+ (White and Kaplan 1972; Ostro and Fondy 1977), and its mitochondrial counterpart the FAD-dependent glycerol-3-phosphate dehydrogenase (mG3PDH, coded by the *Gpd2* gene), which oxidizes G3P back to DHAP and transfers their electrons directly to ubiquinone (Sacktor and Cochran 1957; Estabrook and Sacktor 1958)(Fig. 1A). Several structural and biochemical features make the mG3PDH a unique enzyme. For example, despite G3P oxidation to DHAP is highly exergonic (ΔG = - 37.4 kJ/mol) (Nicholls and Ferguson 2002; Yeh et al. 2008; Mráček et al. 2013), the peripheral location of mG3PDH at the outer leaflet of the inner mitochondrial membrane make it unlikely to conserve the energy of electron transfer as protonmotive force (pmf) (Klingenberg 1970; Yeh et al. 2008). Also, given that mG3PDH transfers the electrons from G3P to ubiquinone this circumvents the reactions of the tricarboxylic acid (TCA) cycle and complex I, making it a suitable alternative path to maintain respiration even in conditions of limited TCA cycle and complex I activities (Chance and Sacktor 1958; Estabrook and Sacktor 1958; Nicholls and Ferguson 2002; Mráček et al. 2013).

**Fig. 1:**
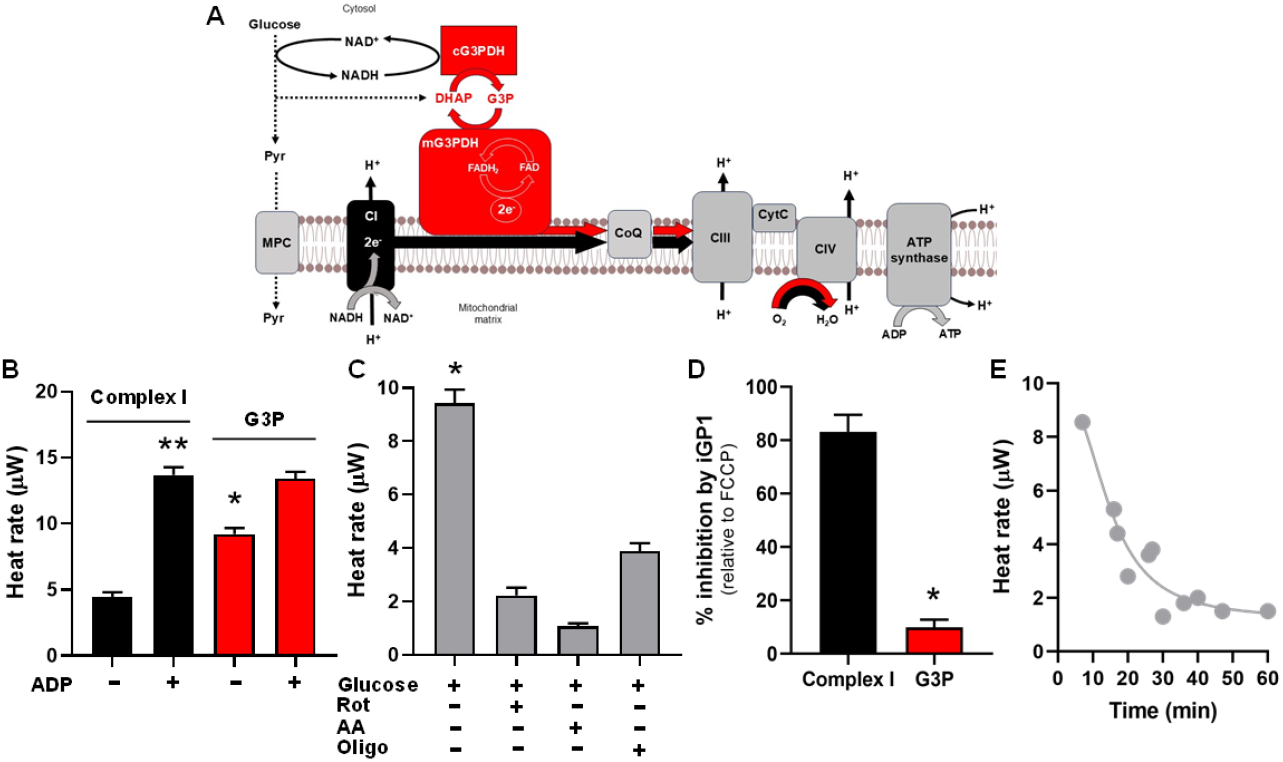
GPSh activity sustains glucose-dependent thermogenesis in *Drosophila* flight muscle. (A) Complex I and mG3PDH heat rate (*µ*W) in permeabilized *D. melanogaster* flight muscles. Bars represent mean ± SEM of calorimetric measurements of 6 fly thoraxes in RB supplemented with: 10 mM pyruvate and 2 mM malate (Complex I) in the absence or in the presence of 5 mM ADP; 20 mM glycerol-3-phosphate (G3P), in the absence or in the presence of 5 mM ADP. ^*^ p < 0.005 relative to complex I without ADP; ^*^ p < 0.0001 relative to complex I without ADP. (B) Glycolytic heat rate (*µ*W) of 6 intact *D. melanogaster* flight muscles in RB supplemented with 5 mM glucose and several OXPHOS inhibitors: 0,5 *µ*M rotenone (Rot), 2.5 *µ*M antimycin a (AA), 1 *µ*g/mL oligomycin (Oligo). ^*^ p < 0.0001 relative to all OXPHOS inhibitors. (C) Inhibitory effect of 80 *µ*M iGP1 on *D. melanogaster* flight muscles determined with 10 mM pyruvate + 2 mM malate + 5 mM ADP + 1 *µ*M FCCP (complex I, black bar) or 0.5 *µ*M rotenone and 20 mM G3P + 5 mM ADP + 1 *µ*M FCCP (G3P, red bar). ^*^ p < 0.05 relative to complex I. Data are expressed as mean ± SEM of at least three different experiments. (D) Inhibitory effect of iGP1 on glycolytic heat production on intact *D. melanogaster* flight muscle. Points represent mean ± SEM of calorimetric measurements of 6 fly thoraxes in RB supplemented with 5 mM glucose and 80 *µ*M iGP-1 along time. Bars represent the mean ± SEM of at least three different experiments.

Insects sustain flight’s extreme energy demands with high metabolic rates, yet their flight muscles are less mechanically efficient (defined as the ratio of mechanical power output to total metabolic power input) than those of birds and mammals (Sohal et al. 1972; Taylor and Heglund 1982; Harrison and Roberts 2000; Mesquita et al. 2021). Interspecific comparisons indicate that efficiency scales positively with body size: while insect muscles convert only about 10% of chemical energy into useful work, avian flight muscles typically achieve 15–25%, and human skeletal muscles reach levels near 70% (Ellington 1985). Interestingly, the relationship between body size and thermogenic ability follows the opposite trend. As size increases, the capacity for internally generated heat declines, yet many insect species remain partially or fully endothermic during flight. Indeed, numerous flying insects maintain thoracic homeothermy, while cursorial species are generally ectothermic and poikilothermic (Harrison and Roberts 2000). In these endothermic fliers, the rapid electron flux through mitochondrial respiratory chains not only fuels ATP production but also releases considerable heat. This coupling of elevated respiration to thermogenic output represents an essential aspect of flight energetics in small-bodied species. Because of their intrinsically low mechanical efficiency, insects compensate by greatly enlarging their mitochondrial oxidative capacity. A striking demonstration of this metabolic adaptation is seen in bumblebees, which activate preflight thermogenesis through preferential oxidation of G3P under cold conditions (Masson et al. 2017).

The G3P oxidation pathway, catalyzed by mG3PDH, is less tightly coupled to ATP synthesis than standard NADH-dependent oxidation (Soares et al. 2015; Masson et al. 2017), allowing more of the redox energy to dissipate as heat. This partial energy non-coupling effectively warms the flight muscles before take□off, ensuring sustained performance even at low ambient temperatures (Soares et al. 2015; Masson et al. 2017). Genetic studies in the fruit fly *Drosophila melanogaster* revealed key physiological and metabolic roles of GPSh activity. Loss-of-function mutants in the *Gpd1* gene manifest profound phenotypes — from flight incapacity and altered mitochondrial ultrastructure to impaired mitochondrial G3P oxidation, indicating critical disruption of energy supply in flight muscles (O’Brien and Shimada 1974; Merritt et al. 2006). Metabolomic profiling reveals compensatory changes in TCA cycle intermediates in maternal-zygotic *Gpd1* mutants, hinting at metabolic rewiring in response to shuttle impairment (Rai et al. 2022). The mutants also show defects in fat metabolism, developmental delays, and shortened lifespan, emphasizing the shuttle’s multifaceted role in energy production, redox balance, and organismal fitness (Merritt et al. 2006; Rai et al. 2022).

We hypothesized that GPSh activity would represent a key mechanism not only for sustaining glycolysis and OXPHOS, but also to mediate heat production. To accomplish this, here we assessed the impacts of pharmacological mG3PDH inhibition on glycolytic activity, mitochondrial G3P oxidation and thermogenesis in *Drosophila* flight muscles. In this sense, we overcame the formidable challenges imposed by the high metabolic rate and diminutive scale of flight muscle samples, by using the specialized chip calorimetry developed by the Freiberg team which enables precise, oxygenated measurements of heat output under physiological conditions. This technology, previously validated in similarly high-activity tissues like brown adipose tissue (Lerchner et al. 2024), ensures reliable thermal recordings while preserving tissue metabolic integrity.

## Material and methods

### Animal stock

*Drosophila melanogaster w*^*1118*^ young males were reared in a B.O.D. incubator at 25×, 12:12 light/dark cycle, 60% relative humidity on a normocaloric diet as culture medium that consists of: 2.83% (w/v) molasses, 157.27 mM glucose, 0.94% (w/v) wheat germ, 0.94% (w/v) soy flour, 41.53 mM sucrose, 1.41% (w/v) cornmeal, 1.08% (w/v) agar, 3.33% (w/v) baker yeast, 9.86 mM methylparaben, 156.2 mM propionic acid, 93% (v/v) distilled H_2_O.

### Sample preparation

Flies aged 1 to 5 days post-eclosion were used in all experiments. Initially, flies were transferred from their culture vials to empty vials devoid of medium. Anesthesia was induced via chill-coma by placing them at 4°C for 2 minutes. Subsequently, the flies were transferred to a Petri dish on an ice bucket for sorting and dissection. Up to 18 individuals were retained on the cold dry petri dish and were dissected in ice-cold respiration buffer (RB, 120 mM KCl, 5 mM KH_2_PO_4_, 3 mM HEPES, 1 mM EGTA, 1,5 mM MgCl_2_, 0.1% w/v bovine serum albumin fatty acid free, pH 7,4) using a scalpel and tweezers. The head, abdomen, wings, and legs were carefully removed from the thorax, and a single incision was made in the isolated thorax using the tip of the scalpel blade.

### Permeabilization

For calorimetry assays, isolated thoraces of up to six individuals were transferred to a Corning PYREX 9 well Spot Plates on an ice bucket and containing 1 mL of ice-cold BIOPS (2.77 mM CaK_2_EGTA, 5.77 mM K_2_EGTA, 5.77 mM Na_2_ATP, 6.56 mM MgCl_2_ 6.H_2_O, 20 mM Taurine, 15 mM NaCl, 20 mM Imidazole, 0.5 mM DTT, 50 mM MES hydrate, pH 7,1) with 5 mg/mL saponin and washed for 30 minutes under gentle stirring at 60 rpm. Following chemical permeabilization, the tissues were carefully transferred using a brush to a separate well containing 1 mL of ice-cold RB and washed for further 10 minutes under the same stirring conditions. Finally, the thoraces were transferred to the calorimeter’s sample holder for further analysis.

### Sample treatment

When sample treatment with specific compounds was required, the prepared thoraxes were incubated outside of the calorimeter for 10 min in the required compound-containing medium, as listed: (a) RB with 10 mM pyruvate and 2 mM malate, (b) RB with 5 mM glucose, (c) RB with 5 mM glucose and 0.5 *µ*M rotenone, (d) RB with 5 mM glucose and 2.5 *µ*M antimycin a, (e) RB with 5 mM glucose and 1 *µ*g/mL oligomycin, (f) RB with 5 mM glucose and 80 *µ*M iGP-1, (g) RB with 10 mM pyruvate, and 2 mM malate and 10 mM ADP, (h) RB with 10 mM pyruvate, and 2 mM malate, 10 mM ADP and 1 *µ*M FCCP, (i) RB with 20 mM G3P, RB with 20 mM G3P and 10 mM ADP, (j) RB with 20 mM G3P 10 mM ADP and 2 *µ*g/mL oligomycin. The measuring channel was filled with the same medium prior to the start of the measurement.

### Calorimetry

To measure heat rates of *D. melanogaster* flight muscle, a chip calorimeter specifically designed for heat rate measurements on small biological samples was used. The calorimeter has already proven to be excellent for calorimetric investigations on brown adipose tissue and other small biological material (Lerchner et al. 2024). Briefly, samples are transported through a medium-filled measurement channel into the heat power detector by a magneto-mechanically driven polymer fiber. The heat power detector consists of a thermopile silicon chip that converts the heat flow dissipated by the sample into a heat power proportional voltage. The heat flow is measured during a constant medium flow through the channel to ensure sufficient oxygenation of the samples. As concluded from previous calorimetric studies on brown adipose tissue using this calorimeter, a complete oxygen atmosphere is required around the heat power detector to avoid anoxic cores within the samples. The measurement procedure was as follows: After filling the measuring channel with the required medium and inserting the transport fiber, three sets of up to six fly thoraxes each were fixed to separate positions of the transport fiber. The sets of thoraxes were then positioned sequentially in the detector and measured under constant medium flow at typically 20 *µ*L.min^-1^.

### Respirometry

Oroboros oxygraph O2K was used to measure the O_2_ consumption rate in mechanically permeabilized *D. melanogaster* thorax as previously described (Gaviraghi et al. 2021). Briefly, four thoraces were dissected and transferred into the oxygraph chamber containing 2 mL of RB at 25ºC, and after closing the chamber the stirrer speed was set to 900 rpm for 7 minutes and 30 seconds. Then the speed was set to 750 rpm. Mitochondrial respiration modulators were added in the chamber with hamilton syringes. Following mechanical permeabilization, the experimental protocol to measure the contribution of G3P to respiratory rates, rotenone (0.5 *µ*M), G3P (20 mM), ADP (5 mM), and iGP1 (80 *µ*M) were added sequentially in a different chamber. In both experiments antimycin A (1 *µ*g/mL) was the last drug to be added to inhibit the ETS.

### Data and statistics

Data in graphs were presented as mean ± standard error of the mean (SEM) of values for each condition. Student’s t test was performed for pairwise comparisons, while multiple comparisons between groups were carried out with one-way ANOVA and *a posteriori* Tukey’s multiple comparisons test. Differences with p < 0.05 were considered significant. Graphs were prepared using the GraphPad Prism software version 8.0 for Windows (GraphPad Software, USA).

## Results

### Mitochondrial glycerol 3 phosphate oxidation is highly thermogenic and contributes to glycolytic heat production

The extent of energy released from the ETS that is dissipated as heat or conserved in the proton gradient can be approximated by comparing proton leak heat rates across substrates. To accomplish this, we make use of chip calorimetry to assess the heat production on both chemically permeabilized flight muscles using complex I substrates or G3P, and whole intact flight muscles using glucose (Lerchner et al. 2024). We observed that in the absence of ADP permeabilized flight muscles, heat production rates by complex I substrates are remarkably lower compared to G3P (4.4 ± 0.4 *vs*. 9.2 ± 0.5 *µ*W, respectively)(Fig. 1B). Interestingly, in the presence of ADP the heat rates generated by complex I substrates or G3P were essentially the same (13.7 ± 0.6 *vs*. 13.4 ± 0.5 *µ*W, respectively)(Fig. 1B). We next investigated the contribution of glucose metabolism to heat production on whole intact flight muscles. The aim was to determine the contribution of glycolysis and OXPHOS on flight muscle heat production when using glucose as a substrate. Figure 1C shows that OXPHOS represents the main contributor to flight muscle thermogenesis as the glucose-dependent heat rates were significantly reduced in the presence of the complex I inhibitor rotenone (Rot), the complex III inhibitor antimycin a (AA), or the ATP synthase inhibitor oligomycin (oligo)(Fig. 1C). The residual thermogenic activity observed after OXPHOS inhibition likely reflects heat production from other glucose-dependent pathways, such as the pentose phosphate pathway and mitochondrial proton leak. To pharmacologically probe the G3P pathway of electron transfer, we used the potent and specific inhibitor of mG3PDH iGP1 (Orr et al. 2014). Confirming the inhibitor specificity, addition of 80 *µ*M iGP1 caused just slight effects (<20%) on the uncoupled respiration sustained by complex I substrates. However, iGP1 strongly reduced uncoupled respiratory rates sustained by G3P oxidation (>98%), demonstrating its potency in *Drosophila* flight muscle (Fig. 1D). To evaluate the role of GPSh in glucose-driven thermogenesis, we measured heat rates in intact flight muscles supplied with glucose and 80 *µ*M iGP1 (Fig. 1E). iGP1 caused a strong, time-dependent inhibition of heat production (∼85%), reaching levels comparable to those under AA treatment. These results indicate that GPSh inhibition disturbs the cytosolic NAD□ redox balance, thereby impairing glycolysis, electron transfer to the ETS, and consequent heat generation. Overall, the data underscores the central role of GPSh in sustaining flight muscle energy metabolism.

## Discussion

In the present work, we investigated how substrate utilization shapes energy coupling and thermogenesis in *Drosophila melanogaster* flight muscle using chip calorimetry. The mG3PDH inhibitor iGP1 almost completely abolished G3P-dependent respiration (>98%), confirming the pathway’s specificity. Calorimetric assays revealed that G3P oxidation generated significantly more heat than complex I substrates, demonstrating its high thermogenic potential. When glucose was supplied to intact flight muscles, OXPHOS inhibitors sharply reduced heat production, while residual rates reflected alternative glucose pathways and proton leak. Inhibition of the GPSh with iGP1 decreased heat generation by ∼85%, highlighting its importance for cytosolic NAD□turnover and glycolytic flux. Together, these results demonstrate that GPSh activity sustains glucose-dependent respiration efficiency and thermogenesis in insect flight muscle.

The heat dissipation rate (J_q_) at a given proton leak rate (J_H+_) depends on the substrate’s oxidation energy (ΔG_ox_) and the number of protons translocated per mole of substrate (r_H+_) from the mitochondrial matrix to the intermembrane space. When proton leak equals proton translocation, no energy is conserved. Assuming minimal reversible entropy contribution (Poe et al. 1967) it reads J_q_≈(ΔG_ox_/r_H+_)J_H+_. The Gibbs energies of oxidation can be derived from the electrochemical potentials of the redox couples (ΔG_ox_= -2FΔE_h_) where F is the Faraday constant, and ΔE_h_ is the difference of the redox potentials of the couples (Nicholls and Ferguson 2002). The redox potentials for NAD□/NADH, DHAP/G3P, and O□/H□O are -320 mV, -190 mV, and +820 mV, respectively, yielding Gibbs energies of approximately -220 kJ/mol for NADH and -187 kJ/mol for G3P oxidation. These values indicate similar oxidation energies for both substrates. Unlike complexes I-IV, which span the inner mitochondrial membrane, mG3PDH is located at the inner membrane’s outer surface (Yeh et al. 2008; Mráček et al. 2013). Although sufficient energy is available when electrons are transferred from G3P to UQ (ΔG = - 37.4 kJ/mol, based on Eh of DHAP/G3P and UQ/UQH2), protons cannot be pumped into the intermembrane space due to mechanistic limitations. Consequently, only six protons are translocated per mole of G3P oxidized, as opposed to ten per mole of NADH. Given the different ΔG_ox_ and proton translocation ratios, the heat rate of proton leakage for G3P should be about 50% higher than for NADH, assuming similar proton leak rates (J_q_(NADH)≈(220/10)J_H+_, J_q_(G3P)≈(187/6)J_H+_). Although the Gibbs energies are approximate, particularly for G3P, and reversible entropy changes may differ for G3P and NADH oxidation, these numerical estimates align with experimental results. Masson et al. discussed the loss of reduction power when FAD is reduced to FADH_2_ due to the oxidation of the glycolytic NADH to NAD^+^ (Masson et al. 2017). However, this is irrelevant in the case of the external introduction of G3P into the system. Therefore, it cannot explain the increased proton leak heat rate that they found in bumblebee flight muscle tissue.

Increased heat production at a given proton flow indicates a loss of respiration efficiency in intact cells. Overall, OXPHOS efficiency can decrease due to either lowered respiration efficiency or impaired coupling between respiration and ATP synthesis. In the flight muscle studied, reduced respiration efficiency seems to be the primary cause of decreased OXPHOS efficiency, while the latter is crucial for heat generation in brown adipose tissue.

Energy coupling in oxidative phosphorylation refers to the precise transfer of energy from electron flow through the ETS to proton pumping, thereby generating a proton motive force (pmf) that drives ATP synthesis. Uncoupling occurs when protons re-enter the mitochondrial matrix bypassing ATP synthase, leading to the dissipation of pmf as heat without ATP production. Acoupling, exemplified by alternative oxidase in plants, involves electron transfer that bypasses key sites of proton transport, releasing energy directly as heat (Elthon and McIntosh 1987; Gnaiger and Group 2020). Based on our results, the contribution of mG3PDH to energy conservation lies between classical coupling and acoupling. Specifically, mG3PDH transfers electrons from G3P oxidation to molecular oxygen via the ETS, where complexes III and IV capture part of this energy as the pmf for ATP synthesis. However, the highly exergonic nature of G3P oxidation (ΔG = − 37.4 kJ/mol) (Nicholls and Ferguson 2002; Yeh et al. 2008; Mráček et al. 2013) and the peripheral localization of mG3PDH on the outer leaflet of the inner mitochondrial membrane (Klingenberg 1970; Yeh et al. 2008) likely prevent efficient conservation of this energy as pmf. Consequently, the G3P oxidation pathway is less tightly coupled to ATP synthesis than NADH-dependent oxidation (Masson et al. 2017), allowing more redox energy to dissipate as heat.

In conclusion, our study shows that mitochondria using G3P as an energy substrate reduces OXPHOS efficiency because the energy released during electron transfer from G3P to ubiquinone is not used to generate the pmf but is instead dissipated as heat. The close association of G3P oxidation with glycolysis in *D. melanogaster* flight muscle indicates high activity of the GPSh, which underlies the thermogenic significance of G3P oxidation in this tissue.

## Data availability

The dataset generated during the current study is available from the corresponding authors on reasonable request.

## Declarations

Conflict of interest: The authors have no conflicts of interest to declare that are relevant to the content of this article.

## Acknowledgements

We are grateful for the valuable comments and criticisms raised by the editor and the reviewers on the original version of the manuscript.

## Author contributions

J.L.: Conceptualization, Supervision, Formal Analysis, Writing – Original Draft, Review. Investigation, Methodology, Data Curation, Writing – Review & Editing. Y.A.R.: Investigation, Data Collection, Formal Analysis, Writing – Original Draft, Review. T.L.: Formal Writing – Original Draft, Investigation, Data Collection, Data Analysis. M.F.O.: Supervision, Critical Review, Final Approval. Conceptualization, Funding Acquisition, Project Administration, Supervision, Writing – Review & Editing, Final Approval.

## References

Chance B, Sacktor B (1958) Respiratory metabolism of insect flight muscle. II. Kinetics of respiratory enzymes in flight muscle sarcosomes. Arch Biochem Biophys 76: 509–531. 10.1016/0003-9861(58)90176-0

Ellington CP (1985) Power and efficiency of insect flight muscle. J Exp Biol 115: 293–304. 10.1242/jeb.115.1.293

Elthon TE, McIntosh L (1987) Identification of the alternative terminal oxidase of higher plant mitochondria. Proc Natl Acad Sci U S A 84: 8399–8403. 10.1073/pnas.84.23.8399

Estabrook RW, Sacktor B (1958) alpha-Glycerophosphate oxidase of flight muscle mitochondria. J Biol Chem 233: 1014–1019

Gaviraghi A, Aveiro Y, Carvalho SS, et al (2021) Mechanical Permeabilization as a New Method for Assessment of Mitochondrial Function in Insect Tissues. Methods Mol Biol Clifton NJ 2276: 67–85. 10.1007/978-1-0716-1266-8_5

Gnaiger E, Group MT (2020) Mitochondrial physiology. Bioenerg Commun 2020: 1–1. 10.26124/bec:2020-0001.v1

Gonçalves RLS, Machado ACL, Paiva-Silva GO, et al (2009) Blood-feeding induces reversible functional changes in flight muscle mitochondria of Aedes aegypti mosquito. PloS One 4: e7854. 10.1371/journal.pone.0007854

Harrison JF, Roberts SP (2000) Flight respiration and energetics. Annu Rev Physiol 62: 179–205. 10.1146/annurev.physiol.62.1.179

Klingenberg M (1970) Localization of the glycerol-phosphate dehydrogenase in the outer phase of the mitochondrial inner membrane. Eur J Biochem 13: 247–252. 10.1111/j.1432-1033.1970.tb00924.x

Lerchner J, Hervas LS, Bícego KC, et al (2024) A device for rapid calorimetric measurements on small biological tissue samples. J Therm Anal Calorim 149: 8085–8096. 10.1007/s10973-024-13183-8

Masson SWC, Hedges CP, Devaux JBL, et al (2017) Mitochondrial glycerol 3-phosphate facilitates bumblebee pre-flight thermogenesis. Sci Rep 7: 13107. 10.1038/s41598-017-13454-5

Merritt TJS, Sezgin E, Zhu C-T, Eanes WF (2006) Triglyceride pools, flight and activity variation at the Gpdh locus in Drosophila melanogaster. Genetics 172: 293–304. 10.1534/genetics.105.047035

Mesquita RD, Gaviraghi A, Gonçalves RLS, et al (2021) Cytochrome c Oxidase at Full Thrust: Regulation and Biological Consequences to Flying Insects. Cells 10: 470. 10.3390/cells10020470

Mráček T, Drahota Z, Houštěk J (2013) The function and the role of the mitochondrial glycerol-3-phosphate dehydrogenase in mammalian tissues. Biochim Biophys Acta 1827: 401–410. 10.1016/j.bbabio.2012.11.014

Nicholls DG, Ferguson SJ (2002) Bioenergetics. Elsevier

O’Brien SJ, Shimada Y (1974) The alpha-glycerophosphate cycle in Drosophila melanogaster. IV. Metabolic, ultrastructural, and adaptive consequences of alphaGpdh-l “null” mutations. J Cell Biol 63: 864–882. 10.1083/jcb.63.3.864

Orr AL, Ashok D, Sarantos MR, et al (2014) Novel inhibitors of mitochondrial sn-glycerol 3-phosphate dehydrogenase. PloS One 9: e89938. 10.1371/journal.pone.0089938

Ostro MJ, Fondy TP (1977) Isolation and characterization of multiple molecular forms of cytosolic NAD-linked glycerol-3-phosphate dehydrogenase from normal and neoplastic rabbit tissues. J Biol Chem 252: 5575–5583

Poe M, Gutfreund H, Estabrook RW (1967) Kinetic studies of temperature changes and oxygen uptake in a differential calorimeter: the heat of oxidation of NADH and succinate. Arch Biochem Biophys 122: 204–211. 10.1016/0003-9861(67)90140-3

Rai M, Carter SM, Shefali SA, et al (2022) The Drosophila melanogaster enzyme glycerol-3-phosphate dehydrogenase 1 is required for oogenesis, embryonic development, and amino acid homeostasis. G3 Bethesda Md 12: jkac115. 10.1093/g3journal/jkac115

Sacktor B, Cochran DG (1958) The respiratory metabolism of insect flight muscle. I. Manometric studies of oxidation and concomitant phosphorylation with sarcosomes. Arch Biochem Biophys 74: 266–276. 10.1016/0003-9861(58)90219-4

Sacktor B, Cochran DG (1957) Dihydroxyacetone phosphate, the product of alpha-glycerophosphate oxidation by insect flight muscle mitochondria. Biochim Biophys Acta 26: 200–201. 10.1016/0006-3002(57)90072-0

Soares JBRC, Gaviraghi A, Oliveira MF (2015) Mitochondrial physiology in the major arbovirus vector Aedes aegypti: substrate preferences and sexual differences define respiratory capacity and superoxide production. PloS One 10: e0120600. 10.1371/journal.pone.0120600

Sohal RS, McCarthy JL, Allison VF (1972) The formation of ‘giant’ mitochondria in the fibrillar flight muscles of the house fly, Musca domestica L. J Ultrastruct Res 39: 484–495. 10.1016/S0022-5320(72)90115-3

Suarez RK, Staples JF, Lighton JRB, Mathieu-Costello O (2000) Mitochondrial Function in Flying Honeybees (Apis Mellifera): Respiratory Chain Enzymes and Electron Flow from Complex III to Oxygen. J Exp Biol 203: 905–911. 10.1242/jeb.203.5.905

Taylor CR, Heglund NC (1982) Energetics and Mechanics of Terrestrial Locomotion. Annu Rev Physiol 44: 97–107. 10.1146/annurev.ph.44.030182.000525

White HB, Kaplan NO (1972) Separate physiological roles for two isozymes of pyridine nucleotide-linked glycerol-3-phosphate dehydrogenase in chicken. J Mol Evol 1: 158–172. 10.1007/BF01659162

Yeh JI, Chinte U, Du S (2008) Structure of glycerol-3-phosphate dehydrogenase, an essential monotopic membrane enzyme involved in respiration and metabolism. Proc Natl Acad Sci U S A 105: 3280–3285. 10.1073/pnas.0712331105

